# Principles for models of neural information processing

**DOI:** 10.1101/129114

**Authors:** Kendrick N. Kay, Kevin S. Weiner

## Abstract

The goal of cognitive neuroscience is to understand how mental operations are performed by the brain. Given the complexity of the brain, this is a challenging endeavor that requires the development of formal models. Here, we provide a perspective on models of neural information processing in cognitive neuroscience. We define what these models are, explain why they are useful, and specify criteria for evaluating models. We also highlight the difference between functional and mechanistic models, and call attention to the value that neuroanatomy has for understanding brain function. Based on the principles we propose, we proceed to evaluate the merit of recently touted deep neural network models. We contend that these models are promising, but substantial work is necessary to (i) clarify what type of explanation these models provide, (ii) determine what specific effects they accurately explain, and (iii) improve our understanding of how they work.

## Introduction

There has been a recent surge of excitement in deep neural networks for neuroscience (Kriegeskorte, 2015; Yamins and DiCarlo, 2016). Major advances in training deep neural networks were achieved by the artificial intelligence and computer vision communities, and these networks now achieve unprecedented performance levels on certain computer vision tasks such as visual object recognition (Krizhevsky et al., 2012). Following these developments, neuroscientists studying the visual system have shown that responses of units in deep neural networks correlate strongly with experimentally measured responses in the primate visual system (e.g., (Agrawal et al., 2014; Cadieu et al., 2014; Eickenberg et al., 2016; Güçlü and van Gerven, 2015a; Khaligh-Razavi and Kriegeskorte, 2014; Kubilius et al., 2016; Yamins et al., 2014)). Due to these correspondences and similarities in architecture between the artificial and biological networks, deep neural networks have been touted as excellent models of biological neural systems.

In this paper, we use the excitement elicited by deep neural networks as an opportunity to think carefully and critically about models of brain function. We step back and consider the broad endeavor of developing models in cognitive neuroscience (Sections 1–2) and provide an assessment of *why* we should develop such models (Sections 3–4). We then highlight the important distinction between functional and mechanistic models (Section 5) and propose specific criteria for evaluating models (Section 6). We end by using the principles we propose to evaluate the merit of deep neural network models (Section 7). While we write this paper as a *Comments and Controversies* article, some of the points in this paper may be uncontroversial, especially to practitioners of model-based neuroscience. However, we think integrating points into a single document is useful, and we believe this consolidated perspective will be especially useful for those who are interested in understanding modeling, but who do not necessarily engage in it.

## 1. What is cognitive neuroscience?

Before reasoning about models in cognitive neuroscience, we must first define these terms. Gazzaniga, Ivry, and Magnun define ‘cognitive neuroscience’ as

> “the question of understanding how the functions of the physical brain can yield the thoughts and ideas of an intangible mind” (Gazzaniga et al., 2014).

It is widely accepted that “thoughts and ideas of an intangible mind,” or mental operations more generally, can be viewed as information-processing operations: for example, the brain represents sensory information, stores sensory information, reasons about this information, and uses information to guide motor behavior. Thus, the brain can be characterized as an organ that mediates interactions between an organism and its environment, accepting incoming sensory information and delivering outgoing motor information. With this definition in mind, the broad question in cognitive neuroscience is, *how do the structure (anatomy) and function (physiology) of neurons and their connections enable the brain to carry out information-processing operations?*

At a coarse level, we already know *what* the brain does, that is, what the information-processing operations are. To use an example from visual neuroscience (DiCarlo and Cox, 2007), we know that one information-processing operation performed by the brain is to take complex spatiotemporal patterns of light impinging on the retina and to use this information to decide what the source of these inputs are (e.g. what type of object is present in the environment). Or, to use an example from social neuroscience (Kubota et al., 2012), we know that one information-processing operation performed by the human brain is to form stereotypes about other humans and use these stereotypes to influence future behavior. But without further work, we do not know *how* the brain performs these operations. The beauty and challenge of neuroscience is that we have tools for measuring in detail the inner workings of the physical organ that performs these operations. Thus, we can move from ‘what does the brain do?’ to ‘how does this particular neuron, or population of neurons, contribute to the functions performed by the brain?’.

## 2. What is a model?

A small but growing number of researchers are using model-based approaches to tackle questions in cognitive neuroscience (e.g., (Brouwer and Heeger, 2013; Forstmann et al., 2011; Huth et al., 2012; Kay and Yeatman, 2017; O’Doherty et al., 2007; Santoro et al., 2014; Sprague and Serences, 2013)). We propose a simple, general definition of ‘model’: a model is a description of a system. In neuroscience, a model would describe how the nervous system is physically structured and/or how its activity changes dynamically over time. In the specific field of cognitive neuroscience, a model would describe how the components of the brain accomplish behaviorally relevant information-processing tasks, such as decision-making or motor control. For example, we might ask, for a given brain region, what stimulus, cognitive, or motor operations are performed by neural activity in that region?

Given the broadness of our proposed definition, nearly any neuroscience result could be viewed as providing a model. However, models vary substantially in how precise and quantitative they are. For example, models can be qualitative, conceptual, and vague about assumptions (e.g., a description in an introductory textbook, or a ‘word’ model that involves ill-defined jargon), or models can be quantitative, mathematical, and explicit about assumptions (e.g., a formal implementation of a model in computer code). Models can depend on concepts and labels derived from our own cognitive abilities (e.g., oracle models that involve manually labeling complex audiovisual stimuli (Huth et al., 2012)), or models can provide explicit specification of concepts and labels independent of a human observer (e.g., a computational implementation of stimulus category (Kay and Yeatman, 2017)). Models can describe systems at coarse levels of detail (e.g., overall activity in a brain region) or at fine levels of detail (e.g., ion channels). As cognitive neuroscientists, we all attempt to describe how the brain performs information processing, and so technically we are all ‘modelers’. Of course, in practice, when we use the term ‘model’, we are typically referring to descriptions that have been made precise and quantitative, and we adopt this usage for the rest of this paper.

## 3. Models make falsifiable claims

Models perform real scientific work, and are not simply *ad hoc* appendages to an experimental study (though in some cases they can be). Rather, models make substantive falsifiable claims and can progressively improve in sophistication and detail. Consider the following simple experiment (Figure 1, left). We ask a human observer to direct her eyes towards a small dot at the center of a blank display. The small dot changes color periodically and we instruct the observer to press a button when the color changes. Meanwhile, we place a stimulus (e.g. a checkerboard) on the display, and move this stimulus to a variety of different positions. As we manipulate the stimulus, we record neural activity in the observer’s occipital cortex using some technology (e.g. fMRI). We discover that there is an increase in activity when the stimulus is present on the display and that there is some variation in activity levels as a function of stimulus position.

**Figure 1.**
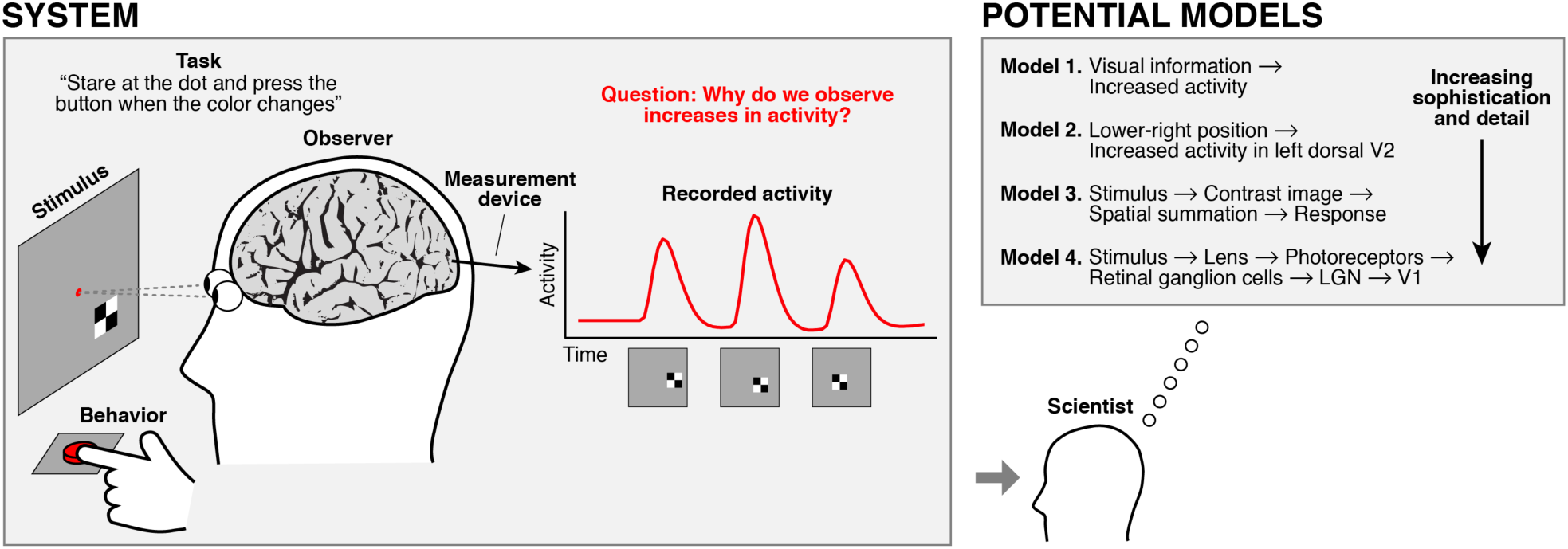
Models describe systems at various levels of sophistication and detail. A typical cognitive neuroscience experiment consists of a stimulus, task, observer, behavior, measurement device, and recorded activity (*left*). A scientist attempts to develop a model of the system, that is, a description of the events that are occurring in the system (*right*). Of particular interest is to characterize why specific levels of neural activity are observed. A variety of different models can be proposed, ranging in sophistication and detail.

In this example, the system consists of the stimulus, task, observer, behavior, measurement device, and recorded activity. Our goal, as scientists, is to describe this system and, in particular, to describe why the increases in neural activity occur. There are many possible descriptions, or models, that we could propose (Figure 1, right). For example, let us consider four potential models:

> *Model 1.* There is visual information on the display (it is not blank). That is why occipital cortex shows increased neural activity.
>
> *Model 2.* There is a point-to-point mapping between positions on the display and positions on cortex (Engel et al., 1997). That is why neural activity at a given cortical position increases for some stimulus positions but not others.
>
> *Model 3.* Spatial extent is one property of a visual stimulus. For a given cortical position, this property is represented through a mathematical operation that takes the spatial extent of the stimulus and performs a weighted sum using a Gaussian function to generate the activity level (Dumoulin and Wandell, 2008). Thus, neural activity levels are what they are because occipital cortex performs this operation.
>
> *Model 4.* Light reflected from the display enters the eye, is refracted by the lens, and is focused onto the retina. Photoreceptors in the retina transduce light energy into electrical voltages. These voltages are communicated by different types of cells to retinal ganglion cells, which send action potentials to the LGN. In turn, neurons in the LGN send action potentials to primary visual cortex. At each stage in this process, sensitivity is local (e.g., photoreceptors are sensitive to light from a restricted region of the visual field, neurons in the LGN receive input from a specific collection of neighboring retinal ganglion cells, etc.). The net result of these processing stages can be summarized by any of the earlier three models.

Although the above models vary widely in sophistication and detail (and we could go into even further detail with respect to molecular mechanisms), all of the models describe the system under consideration and make substantive falsifiable claims. Each model posits certain variables as being causally related to the observed neural activity and implicitly excludes other variables. The claim is that the visual stimulus matters to the neural activity, but that for example, the auditory background noise that happened to be present during the experiment, the motor behavior, and the internal cognitive state of the observer do not. With additional experimental measurements, we can test whether the models are indeed sufficient or whether modifications to the models are necessary. If we find that variables such as auditory stimulation or cognitive state affect the observed activity, these variables must be included to achieve a complete description of the system.

The examples provided above, like many studies in cognitive neuroscience, characterize neural activity in specific brain regions. This approach assumes we have already accurately identified the relevant brain regions in a given observer, which can be a challenging endeavor (Benson et al., 2012; Frost and Goebel, 2012; Glasser et al., 2016; Gordon et al., 2016; Sabuncu et al., 2010; D. Wang et al., 2015; L. Wang et al., 2014; Weiner and Grill-Spector, 2012). We stress the importance of careful localization of brain regions before developing models of their function. An increasing number of researchers are developing quantitative models of where distinct regions and networks are located within the brain (Haxby et al., 2011; Huth et al., 2016; Nelson et al., 2010; Yeo et al., 2011). Interestingly, locations of regions and networks in cortex do not appear to be random and are instead very predictable. Recent research indicates this predictability may stem from several types of neurobiological substrates. For example, cortical folding (Benson et al., 2012; Weiner et al., 2014), white matter (Saygin et al., 2011; Yeatman et al., 2014), cytoarchitectonics (Rosenke et al., 2017; Weiner et al., 2016a), and myelination (Glasser et al., 2016) can all contribute to predicting the locations of functional regions.

## 4. Why are models useful?

Developing precise and quantitative descriptions of how the brain performs information processing takes effort. In our view, models provide three main benefits: summary, explanation, and prediction. We describe these benefits below, and refer the reader to a concrete example taken from previous work (Figure 2).

**Figure 2.**
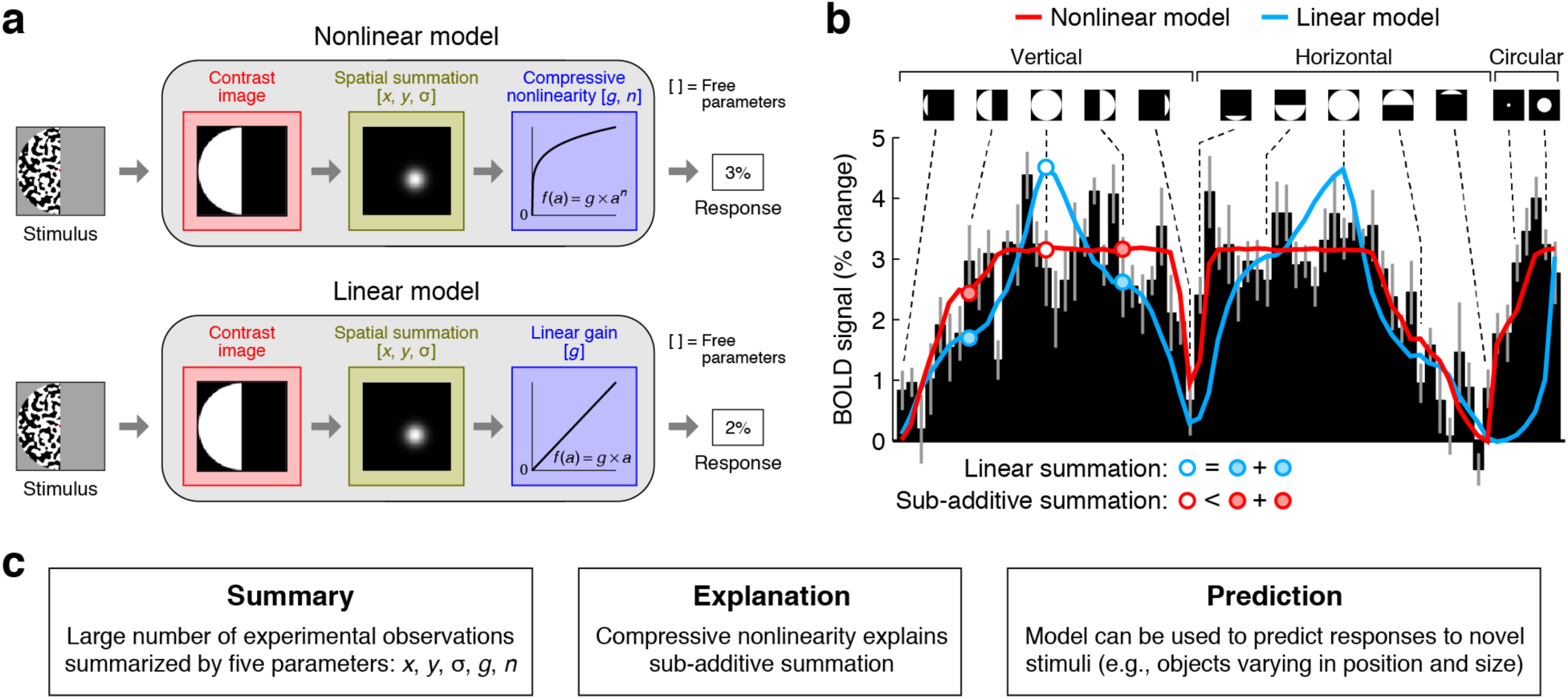
A concrete example of how models provide summary, explanation, and prediction. Figure adapted from (Kay et al., 2013a). *a*, Two potential models of how spatial extent of visual stimuli relates to neural responses. The nonlinear model starts with a contrast image representing stimulus location, computes a weighted sum of this contrast image using a 2D Gaussian, and applies a compressive nonlinearity. The linear model is identical except that the compressive nonlinearity is removed, leaving a linear gain. *b*, Data and model predictions for a voxel in visual area TO-1. Black bars indicate measured BOLD responses to different stimulus locations (depicted by small icons). Leave-one-stimulus-out cross-validation was used to fit the models, and thick lines indicate model predictions. An effect of interest is whether the response to a full stimulus (open dots) is equal to the sum of the responses to two partial stimuli (filled dots). The data support sub-additive summation, which is captured by the nonlinear model. *c*, Three functions performed in this modeling example. (1) The nonlinear model *summarizes* the large set of noisy measurements using just five parameters. (2) The removal of the compressive nonlinearity leads to linear summation, which does not match the data; thus, the compressive nonlinearity is necessary for, and *explains*, sub-additive summation. (3) The nonlinear model *predicts* responses to novel stimuli. For example, the nonlinear model predicts specific levels of tolerance in responses to objects varying in position and size, and experimental measurements have confirmed this prediction (Kay et al., 2013a).

### Summary

Neural measurements are complex and noisy, and there is no limit to the number of experimental variations that one can investigate. Models can provide compact summaries of the information processing that a neural system is performing. Thus, a major benefit of a model is that one can make inferences on a focused set of parameters that summarize the data, instead of attempting to interpret a large number of noisy individual data points. Parameters derived in this way can then be compared across brain areas (e.g. (Kay et al., 2013b)) or subject populations (e.g. (Schwarzkopf et al., 2014)).

### Explanation

Models posit that specific variables relate to neural activity. As such, models provide *explanations* of measurements of the brain. For example, suppose we find that a neuron is highly active when a clip of rock music is played but is only weakly active when a speech clip is played. Why does this occur? One model could be that the neuron computes overall sound intensity, and the reason we observe weak activity for the speech clip is that it has low sound intensity. Alternatively, there are other candidate models that might explain the phenomenon (e.g., selectivity for guitar tones, selectivity for speech, variations in attentional engagement). With appropriate experimental measurements, we can adjudicate different models and decide which model is most accurate (Naselaris and Kay, 2015).

### Prediction

There are several different senses in which models provide predictive power. One sense comes from cross-validation (Hastie et al., 2001), a procedure that is commonly used in model-based studies. In cross-validation, the researcher sets aside some testing data, fits the parameters of a model on the remaining training data, and then assesses how well the fitted model predicts the testing data. The testing data could reflect distinct trials of the same experimental conditions found in the training data, in which case this demonstrates limited predictive power.

Alternatively, the testing data could reflect completely novel experimental conditions, which demonstrates stronger predictive power.

A different sense in which models provide predictive power is if a model developed in one study is able to predict the results of a new study that does not involve exactly the same subjects, stimuli, and task design used in the first study. For example, we have shown that a model developed using simple artificial stimuli and fMRI measurements successfully generalizes to complex naturalistic stimuli (Kay et al., 2013a) and data obtained from a different measurement technique (Winawer et al., 2013). As another example, we have shown that a model that describes structural-functional relationships in one group of subjects can successfully generalize to a new group of subjects (Rosenke et al., 2017; Weiner et al., 2016a).

A third and deep sense in which models provide predictive power is if a model is able to predict the consequences of physical perturbations to the brain. If we had accurate and detailed descriptions of how neural systems in the brain coordinate to perform information-processing operations, we should be able—in principle—to predict, for example, the effects of lesions made in specific brain areas (Gallant et al., 2000), surgical removal of entire brain areas (Weiner et al., 2016b), the effects of enhancement (Salzman et al., 1990) or disruption (Pascual-Leone and Walsh, 2001) of neural activity, or the effects of psychoactive drugs (Rokem and Silver, 2010). These are not easy predictions to make, assuming we are careful to avoid the illusion of predictive power that comes from making “predictions” after looking at the data. A model conjured to explain effects that have already been observed generates “postdictions” and should be treated with skepticism (since the model has been de facto fit and tailored to the data).

## 5. Functional vs. mechanistic models

It is important to distinguish between *functional models* and *mechanistic models* of neural information processing (Albrecht et al., 2002; Carandini, 2012; Carandini and Heeger, 2011). Functional (or computational) models characterize the transformation between input and output performed by a neuron or population of neurons (Wu et al., 2006), reminiscent of the concept of functions in mathematics or programming. Mechanistic (or biophysical or circuit) models characterize the details of the method or mechanism by which a neuron or population of neurons carry out such a transformation (Priebe, 2016). To illustrate, recall Models 1–3 from the previous example. These models are all stimulus-referred (Heeger et al., 1996; Wandell et al., 2015) in the sense that they specify how the stimulus relates to activity in occipital cortex. Thus, the models can be viewed as functional models that characterize the transformation between input (stimulus) and output (neural activity). In contrast, Model 4 concerns not only the stimulus, but also the series of physical events that intervene between the stimulus and neural activity. This model can therefore be viewed as a mechanistic model that characterizes how the brain carries out the transformation described by Models 1–3. There may be multiple possible mechanistic models that are all consistent with a given functional model. Functional models can be rigorously established for a system, even if the underlying mechanisms are not yet known.

Functional and mechanistic models are complementary to one another and should be judged on their own merits. The value of functional models is that they emphasize the outcomes and meaning of neural information processing. The significance of a signal carried by a neuron or population of neurons ultimately lies in what that signal conveys about sensory or motor information for the observer. For example, if an organism encounters a predator, what matters is successful detection of the predator so that motor behavior can be appropriately guided; how that detection is accomplished is of secondary importance. Focusing on mechanisms without addressing sensory or motor significance would generate an incomplete picture of neural information processing. On the other hand, the value of mechanistic models is that they are necessary for a complete understanding of a system. To fully describe a functioning brain, we must specify not only what it and its parts compute, but also how they actually compute it. The presumption—at least under conventional neuroscientific thinking—is that the overall information-processing tasks performed by the brain are ultimately the result of neurons and their interactions, which are governed by basic principles of chemistry and physics. However, therein still lies immense complexity, given the sheer number of neurons, cell types, and connections in the brain.

These points are directly related to David Marr’s well-known levels of analysis where distinctions are made among computational, algorithmic, and implementation levels (Marr, 1982). Slightly generalizing our definition of ‘mechanistic’ to refer to the details of how something is accomplished, we see the algorithmic level serves as a mechanistic model for the computational level and the implementational level serves as a mechanistic model for the algorithmic level. For example, imagine a situation where an organism is attempting to determine the location of a predator from auditory inputs. We can describe the system at a computational level by characterizing the problem that the organism is trying to solve: given auditory inputs, detect the predator and determine the direction of the predator. We can describe the system at an algorithmic level by identifying the specific set of auditory and decision-making algorithms that the brain uses to solve the problem. Or we can describe the system at an implementational level by identifying the specific configurations of neurons and connections that implement those algorithms. Each level provides details as to how the level above is accomplished. Presumably, we should strive to build accurate models at all levels of analysis.

Many studies in cognitive neuroscience develop functional models and ignore anatomical implementation. For example, a researcher might use fMRI to investigate how patterns of population activity represent a stimulus, irrespective of details of how this activity is spatially organized across cortex. Or, a researcher might use electrophysiology to study how individual neurons respond to experimental conditions, irrespective of details of cell types or the circuit that a neuron participates in. Whether this is a fruitful or fruitless approach is open for debate. Although it is not entirely known how anatomical details—from cortical folding patterns down to the level of neuron density, cell types, and ion channels—constrain and shape functional processing (Van Essen, 1997), we believe that anatomy may hold valuable clues to function. If the brain spends so much neurobiological energy organizing structure and function across spatial scales, presumably this orderliness is useful for something. For example, perhaps the specific way that functional properties are clustered in the brain enables faster and more efficient readout of information for a particular task (Grill-Spector and Weiner, 2014). However, we cannot rule out that in some cases, there may be anatomical structures without any particular functional meaning (Horton and Adams, 2005). The path to understanding anatomical implementation will be difficult, as even the heavily studied functional property of orientation selectivity in primary visual cortex has not been resolved in terms of circuit-level mechanisms (Priebe and Ferster, 2008). Nonetheless, a deeper understanding of anatomical implementation is critical, especially if we want to be able to predict the effects of physical perturbations to the brain or make predictions that successfully span vast differences in spatial scale of measurement in neuroscience (Sejnowski et al., 2014).

## 6. What makes a good model?

Thus far, we have addressed what models of neural information processing are, why they are useful, and the distinction between functional and mechanistic models. Now suppose in our daily work, we come across a model put forth by a researcher in the field. How should we evaluate the merit of the model? We propose two criteria, accuracy and understanding.

### Accuracy

The first criterion is accuracy, which refers to how well a given model performs in matching the system under investigation (for example, see Figure 2b). To assess accuracy, we collect experimental data at some spatial and temporal scale, perform proper preparation and binning of those data, and then quantify whether the predictions of a model match the data. Typically, models have free parameters whose values are not known *a priori* and must be set to obtain quantitative predictions. These parameters are usually set based on experimental data, and in such cases, it is crucial to control for overfitting. This can be done by evaluating predictive performance on left-out data (i.e. cross-validation) or by using techniques that penalize goodness-of-fit based on number of free parameters (e.g. Akaike Information Criterion). It is sometimes disparagingly remarked that a model is ‘just fitting the data’—on the contrary, quantitatively matching experimental measurements is exactly what a model ought to do.

Beyond quantifying predictive power for a set of data, we should consider the range and diversity of the experimental manipulations represented by those data. A model should describe how the brain carries out information processing in a broad range of situations, not just the specific situations used in one or a few particular studies (Felsen and Dan, 2005; Kay et al., 2013b). For example, suppose visual sinusoidal gratings are presented to an observer and a model is proposed in which the activity of a neuron is calculated as a weighted sum of the luminance values of the stimulus (Carandini et al., 2005). This model posits that neural activity reflects a specific visual attribute and, by implication, does not reflect other visual, cognitive, or motor attributes. Evidence for the accuracy of the model would be greatly strengthened if we performed a diverse range of experimental tests—for example, using naturalistic visual scenes to deliver luminance stimulation (David et al., 2004), manipulating the internal cognitive state of the observer (McAdams and Reid, 2005), or allowing visual stimulation to occur simultaneously with motor responses—and still found that the same model (with exactly the same parameters) accurately predicts neural activity. By performing stringent tests of a model, we gain confidence in its accuracy.

We comment briefly on the topic of phenomenological models. It is possible to have a model that accurately matches a set of data, but performs no actual explanatory work. Such models (which can also be termed ‘purely descriptive models’) may be useful for comparison purposes, but do not provide neuroscientific insight (Albrecht et al., 2002). For example, suppose we are investigating how neural responses to stimuli change as a function of the cognitive task that a subject is performing (Kay and Yeatman, 2017). We could propose a model that allows each task to induce an additive offset to neural responses, and this model could be fit and evaluated like any other model. However, the model does not make a substantive claim about the specific property of the tasks that is responsible for the additive offsets, and therefore has limited neuroscientific value. (Imagine trying to predict responses for a novel cognitive task—the model would be incapable of doing so because it does not provide any insight into the nature of cognitive tasks.)

### Understanding

The second criterion for the merit of a model is understanding, which refers to how well we, as scientists, grasp the relationship between the components of a given model and the outcomes that the model predicts. Or, in simpler terms, *do we know how the model works?* To illustrate, suppose we observe neural activity is higher in one experimental condition compared to another. A model that describes this system should indicate what property of the first condition leads to increased neural activity. If the model successfully conveys what this property is, we will have *understood* why the effect occurs. In practice, models can be mathematically or algorithmically complex, and it may take effort to determine which specific model component is responsible for a given effect (for an example of how this can be done, see Figure 2).

It is helpful to consider examples where model understanding is poor. Suppose we wish to characterize the relationship between two continuous variables, *x* and *y*. One approach is to characterize *y* as a weighted sum of the outputs of nonlinear basis functions defined on *x* (e.g., the weighted sum of a large number of Gaussian functions). Another approach is to simply characterize *y* as a linear function of *x*. Now suppose the relationship between *x* and *y* is, in fact, linear. Both the complex nonlinear model and the simple linear model are identical in their behavior and equally accurate in matching the data. However, the complex model has less value because it provides less understanding: to understand the model, we would have to expend additional effort analyzing the tuning properties of the basis functions and the weights associated with the basis functions.

As another example, suppose we have two code implementations of a functional model of neural information processing, one set of code being short, concise, and well-documented, the other set of code being long, convoluted, and undocumented. Both sets of code behave identically in their input-output behavior and achieve the same accuracy in matching experimental data. However, the longer code has less value because it provides less understanding: to figure out what model the code implements, we have to pore through and digest computer code. More generally, this example highlights the importance of clarity in descriptions of models. Clarity should exist at the verbal or conceptual level (scientific prose), the mathematical level (equations), and for models that are algorithmically complex, the computational level (code). An accurate model is useless if not clearly described.

What are some practical methods for improving understanding of a model? One is to simply observe the model’s behavior. Observing how a model behaves across different experimental manipulations is useful, even if empirical measurements of those manipulations are not available. For example, a functional model of visual processing could be probed using a variety of different stimulus manipulations, such as changing the orientation of a bar, changing the semantic category of a object, etc. Carefully controlled experimental manipulations help isolate and identify what effects are explained by a given model (Rust and Movshon, 2005). A second method for improving understanding is to manipulate the model and examine the effect on the model’s behavior (Kay et al., 2013b; Nishimoto and Gallant, 2011). If we remove a certain model component or change a certain model parameter, does the model fail to account for the effect of interest? If the model fails, we have learned that the identified component or parameter is critical (for an example, see Figure 2). If the model still works, we have learned that the identified component or parameter is not critical, and we could remove it to obtain a simpler and easier-to-understand model. A third method is to model the model, that is, perform simulations of the model’s behavior and attempt to develop a simpler model that accounts for the observed behavior. For instance, in the previously described example involving variables *x* and *y*, we could take the complex nonlinear model, perform simulations, and eventually realize that a simple linear model can reproduce the model’s behavior.

## 7. The case of deep neural networks

Now that we have covered principles for assessing models of neural information processing, we turn to the specific case of deep neural networks (DNNs). These networks, inspired by properties of biological visual systems (Fukushima, 1980; Serre et al., 2007), consist of multiple layers of processing, where each layer is composed of units that perform relatively simple linear and nonlinear operations on the outputs of previous layers. Connections between units are typically designed such that a convolution is performed in the linear weighting step (same weights are applied at different positions), which parallels the visual system. Parameters of the networks are typically set using supervised learning techniques, optimizing performance on specific tasks such as predicting the object category associated with the visual input (Yamins et al., 2014). Researchers have demonstrated high levels of correlation between activity exhibited by DNN units and measurements of activity in visual cortex in response to naturalistic objects and scenes (Eickenberg et al., 2016; Güçlü and van Gerven, 2015b; Khaligh-Razavi and Kriegeskorte, 2014; Yamins et al., 2014).

Do DNNs have merit as models of biological visual systems? The answer depends on the specific claim that is being made. Suppose the claim is simply that activity in visual cortex reflects a series of processing operations that are performed on visual inputs provided to an observer. This minimal interpretation, that ‘visual cortex is a multi-layer neural network that processes visual inputs’, is a simplistic qualitative model that is neither exciting nor objectionable, but nevertheless counts as a valid model (see Section 2). Presumably, there is a deeper, more substantive claim that one wants to make regarding DNNs, and the merit of this claim will depend heavily on details of the architecture and parameters used in a DNN. Do we wish to adopt the extreme claim that every parameter value in a DNN is critical and every DNN unit corresponds to a specific neuron or neural population in the brain? If not, what is the proposed interpretation?

An important distinction that affects the interpretation of DNNs is whether they are intended as functional or mechanistic models (see Section 5). Suppose DNNs are intended only as functional models of how stimuli (inputs) relate to neural responses (outputs). In this case, there are a number of open questions to address regarding the accuracy of DNNs. Thus far, researchers have examined large-scale datasets involving a diversity of complex naturalistic stimuli and demonstrated general correspondence between artificial and biological responses. However, much work in visual neuroscience has characterized in detail how specific visual areas represent specific stimulus dimensions, such as contrast (Albrecht et al., 2002), spatial extent (Kay et al., 2015), curvature (Brincat and Connor, 2004), color (Horwitz and Hass, 2012), and spatial frequency (Lennie and Movshon, 2005), just to name a few. Do DNNs accurately account for these effects? Furthermore, we should scrutinize the range and diversity of experimental manipulations that have been examined (see Section 6). DNNs provide potential explanations of stimulus-driven activity, but these are ultimately incomplete descriptions given that visual activity is affected by non-stimulus factors, such as attention (Luck et al., 1997), imagery (O’Craven and Kanwisher, 2000), and working memory (Harrison and Tong, 2009).

Suppose instead that DNNs are intended as mechanistic models that not only characterize stimulus-response transformations, but also the way in which the brain accomplishes those transformations (see Section 5). If this is the intention, we again are faced with a number of open questions. What is the proposed mapping between individual units in a layer of a DNN and the neurons in a given brain area? Are DNNs attempting to account for variations in the physical sizes of different visual areas (Dougherty et al., 2003)? Do layer-to-layer connections in a DNN accurately reflect physical connections in biological visual systems, e.g., the spatial extent of V1 neurons that project to a V2 neuron (Sincich et al., 2003)? How can we reconcile DNNs with the existence of bypass routes in corticocortical connections (Felleman and Van Essen, 1991) which violate a strictly hierarchical organization? Can DNNs account for different cell types, the laminar organization of cortex, and the existence of extensive feedback projections?

In addition to accuracy, we should also evaluate our understanding of DNNs. The computational capabilities of DNNs depend critically on the specific parameters used in the models (Coates et al., 2011; Pinto et al., 2009). However, DNNs have many thousands (or even millions) of free parameters, and so understanding DNNs is not an easy task. If we fail to understand DNNs (either because they are too complex or because we do not want to) and treat these models as ‘black boxes’, they perform the function of prediction but fail to perform the functions of summary and explanation (see Section 4). They do not summarize well because there are too many potential parameters that contribute to the description of the system; they do not explain well because it is not clear which specific parts of the models are necessary to explain a given effect.

Overall, we are not claiming that DNNs have no utility, but we are highlighting open questions and limitations that apply to DNNs (as well as other potential models of neural information processing). There is substantial work to be done towards clarifying what type of explanation these models are supposed to provide and determining what specific experimental effects they accurately explain. We need also to improve our understanding of how these models work. There are concrete steps we can take towards improving understanding (as discussed in Section 6). We can observe the models (e.g., inspect responses to controlled stimuli (Eickenberg et al., 2016)), we can manipulate the models (e.g., perturb parameters and examine the consequences (Cichy et al., 2016)), and we can model the models (e.g., perform simulated experiments ‘in silico’ and derive simpler models that achieve the same behavior). The concerns we voice here have guided our own research approach which produces models that explain specific effects and have components that are well understood (e.g., (Kay et al., 2013b; Kay and Yeatman, 2017)).

## Conclusion

We wrote this perspective at a broad level to remove us from the messy, often confusing, details of different measurement methods (e.g., fMRI, EEG/MEG, electrophysiology), different data analysis approaches (e.g., multivariate pattern analysis, representational similarity analysis, voxelwise modeling, functional connectivity), and jargon (e.g., encoding, decoding). Although technical details matter (Naselaris et al., 2011), the goal of this paper is to emphasize the larger point that we should be using measurements of the brain to build models of how neurons and neural populations perform complex information-processing operations. These models should accurately predict what happens under a broad range of experimental manipulations, and we should understand these models through clear description, observation, and manipulation. When we encounter a model in the literature, we should carefully consider questions such as: How well do they account for the data? How extensive are the experimental manipulations? How clear is the link between the components of the proposed model and the observed effects? Is the model attempting to provide a functional or mechanistic explanation?

It is useful to draw inspiration from other domains of science. In chemistry, we know if we mix a certain amount of one chemical with another, we will observe certain outcomes, such as emission of heat. This is because we have working models of the relevant variables (e.g., molecular composition of the chemicals) and how these variables interact. In astronomy, we know if we observe two celestial bodies headed towards each other, we will observe certain outcomes, such as collision or trajectory deviation. This is because we have working models of the relevant variables (e.g., mass, velocity, presence of other nearby bodies) and how these variables interact. In cognitive neuroscience, suppose we developed models that could predict the behavioral and neural outcomes of an arbitrary experiment involving stimuli and task instructions. Such models would predict how fast and accurate an observer will be at the task, what levels of neural activity will be found in different brain areas, and how these neural activity levels relate to the sensory, cognitive, and motor processes involved. Once we achieve such models, we might be able to claim to know how the brain works.

## Acknowledgments

We thank B. Hutchinson, M. Moerel, N. Rust, and J. Winawer for comments on the manuscript. Portions of this work were presented at a symposium held at Vision Sciences Society 2016.

